# TMPRSS6 Cleavage of β-Klotho Modulates FGF19 Signaling

**DOI:** 10.64898/2026.08.26.746010

**Authors:** Matthieu Lepage, Antoine Désilets, Gabriel Lemieux, Michael Desgagné, Pierre-Luc Boudreault, Richard Leduc

## Abstract

Metabolic dysfunction-associated steatotic liver disease (MASLD) is the most prevalent liver disorder worldwide, yet therapeutic options remain limited. TMPRSS6, a liver serine protease best known for its role in iron homeostasis, has recently emerged as a potential therapeutic target for MASLD. However, the molecular mechanisms linking TMPRSS6 to hepatic lipid metabolism remain incompletely understood. To identify novel TMPRSS6 substrates, we performed extracellular proteomic analyses of TMPRSS6-overexpressing cells. Among the proteins identified, β-klotho (KLB), a co-receptor required for FGF19 and FGF21 signaling, emerged as a compelling candidate substrate. We demonstrate that TMPRSS6 interacts with KLB and promotes its proteolytic shedding in a catalytic activity-dependent manner. Functionally, TMPRSS6 reduced full-length KLB abundance at the cell surface and attenuated FGF19-dependent FGFR4 signaling in a heterologous expression system. Together, these findings identify KLB as a novel functional substrate of TMPRSS6, providing a mechanistic framework through which this protease may influence hepatic lipid metabolism. These results provide a rationale for investigating the regulation of KLB and other candidate substrates by TMPRSS6 in physiological models and further support its evaluation as a therapeutic target for MASLD.

## Introduction

Metabolic dysfunction-associated steatotic liver disease (MASLD) is currently the most common liver disease, affecting 38% of the global adult population, with prevalence projected to reach 55% by 2040^1,2^. This condition is characterized by hepatic lipid accumulation that can progress to the more severe stage known as MASH (metabolic dysfunction-associated steatohepatitis), the inflammatory and fibrotic form of the disease that can ultimately lead to cirrhosis^3,4^ and hepatocellular carcinoma^5^.

Lifestyle interventions aiming at weight loss, including physical activity and a healthy diet, are the main treatment option, but long-term compliance and efficacy are often limited for many individuals^6,7^. Few pharmacological treatments exist. To date, two molecules, resmetirom, a selective thyroid hormone receptor-β agonist, and semaglutide, a glucagon-like peptide-1 receptor agonist, have recently received FDA approval for the treatment of MASH^8,9^. However, their long-term efficacy and safety remain under evaluation, and neither therapy is currently approved for patients in earlier stages of MASLD. In this context, developing alternative or complementary therapeutic strategies is important to broaden the range of interventions available for MASLD and MASH.

A potential emerging target for the treatment of MASLD is the transmembrane protease serine 6 (TMPRSS6), also known as matriptase-2. TMPRSS6 belongs to the type II transmembrane serine protease (TTSP) family and is predominantly expressed in hepatocytes, where it is best known for its central role in iron homeostasis through negative regulation of hepcidin via the BMP-SMAD pathway^10–13^. Hepcidin is the key hormone responsible for regulating systemic iron levels by binding to ferroportin and inducing its degradation thereby controlling iron export from enterocytes and macrophages into plasma^14,15^. However, the exact mechanisms by which TMPRSS6 regulates hepcidin expression remain unclear. TMPRSS6 cleaves the BMP co-receptor hemojuvelin (HJV) and other components of the BMP-SMAD pathway^16,17^, but these findings lack *in vivo* validation and evidence suggests that TMPRSS6 may also modulate this pathway through non-proteolytic mechanisms^18,19^.

Beyond its established role in iron homeostasis, accumulating evidence suggests that TMPRSS6 also influences metabolic pathways involved in hepatic lipid metabolism and MASLD pathogenesis. Genetic studies have identified TMPRSS6 variants as risk factors for MASLD^20^ while TMPRSS6 expression is elevated in affected livers^21^. Mechanisms implicated in MASLD pathogenesis intersect with pathways regulated by TMPRSS6. For example, reduced BMP-SMAD signaling promotes MASLD and lipid dysregulation^21^. More directly, TMPRSS6 knockout (KO) mice are resistant to diet-induced obesity and hepatic steatosis^22,23^, and exhibit reduced hepatic inflammation^24,25^. Furthermore, TMPRSS6 downregulation in mouse hepatocytes counteracts steatosis and inflammation, further highlighting its potential role in disease progression^21^. Although these findings identify TMPRSS6 as a novel modulator of lipid metabolism and as a promising therapeutic target for MASLD, the mechanisms underlying this regulation remain incompletely understood. It has been proposed that TMPRSS6-dependent inhibition of the BMP-SMAD pathway suppresses PPARα signaling, thereby contributing to MASLD-MASH progression^21^. Whether this mechanism fully accounts for the metabolic effects of TMPRSS6 remains unclear and additional mechanisms may contribute. Given that TMPRSS6 is a membrane-bound serine protease, identifying novel substrates may provide important insight into its role in hepatic metabolism.

To this end, we performed proteomic analyses to identify novel TMPRSS6 substrates. Among the candidates identified, β-Klotho emerged as a particularly compelling candidate because of its central role in FGF19/FGF21 signaling. Our findings reveal a previously unrecognized link between TMPRSS6 and the FGF19/FGF21-KLB signaling axis and suggest a potential mechanism through which TMPRSS6 may influence hepatic metabolism.

## Results

### TMPRSS6 modulates the extracellular proteome of Hep3B cells

As a transmembrane protease, TMPRSS6 is expected to cleave transmembrane or membrane-anchored proteins, resulting in the release of protein fragments into the extracellular medium. Characterizing the extracellular proteome therefore provides a relevant approach to identify novel TMPRSS6 substrates, as well as downstream secreted effectors regulated by TMPRSS6, such as hepcidin^12,13^. To do so, the hepatocellular carcinoma Hep3B cells were first transfected with either an empty vector (Mock) or TMPRSS6 WT, and the conditioned media were concentrated before mass spectrometry analysis to identify shed or secreted proteins. Identified proteins were then computationally filtered using gene ontology (GO) cellular component annotations to exclude intracellular contaminants, and the resulting extracellular proteome was subjected to differential abundance analysis (Fig. 1A). Several proteins were significantly enriched (n=12) or reduced (n=5) using this procedure, confirming that TMPRSS6 modulates Hep3b extracellular proteome. Furthermore, to distinguish between non-catalytic and catalytic-based mechanisms of regulation, TMPRSS6 WT was also compared to another condition overexpressing the catalytically inactive mutant TMPRSS6-S762A which resulted in 15 significantly enriched proteins and 13 significantly reduced proteins (Fig. 1B).

**Figure 1.**
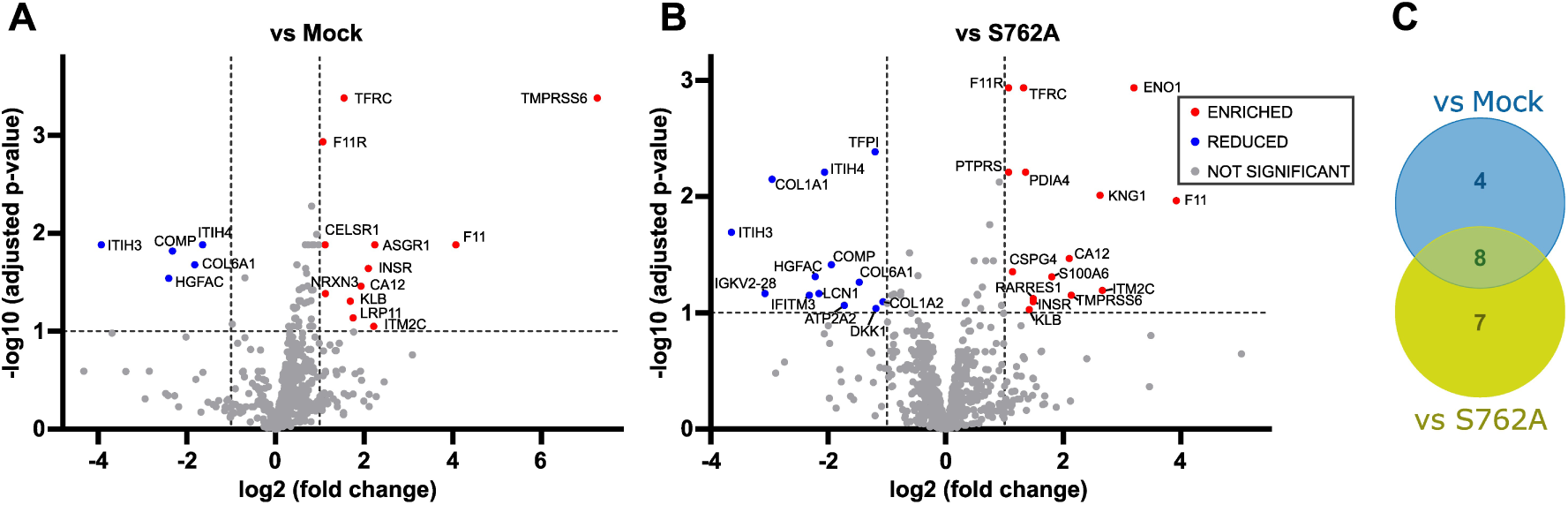
TMPRSS6 modulation of the extracellular proteome. (A-B) Volcano plot showing changes in extracellular protein abundance identified by mass spectrometry in the conditioned medium of Hep3B cells overexpressing TMPRSS6 compared with mock-transfected cells (A) or cells overexpressing the inactive TMPRSS6 S762A (B). Proteins significantly enriched (fold change ≥ 2, adjusted p-value < 0.1) are shown in red, and those decreased (fold change ≤ 2, adjusted p-value < 0.1) are shown in blue. (C) Venn diagram of proteins significantly enriched in the condition overexpressing TMPRSS6 compared to both mock and S762A.

To prioritize the most robust candidate TMPRSS6 substrates, we identified proteins that were consistently enriched in both the TMPRSS6 WT versus Mock and TMPRSS6 WT versus TMPRSS6-S762A comparisons, leading to eight candidates (Fig. 1C, Table 1). Proteins consistently reduced in both comparisons were also identified (Fig. S1, Table S1) as these may represent secreted proteins whose abundance is indirectly regulated by TMPRSS6. Among the eight proteins enriched in both comparisons, seven are expressed in the liver (>1 nTPM) according to the Human Protein Atlas database (https://www.proteinatlas.org/)^26^. Six of these are located at the plasma membrane including two previously reported TMPRSS6 substrates; TMPRSS6 itself, which undergoes autoproteolytic cleavage^27,28^, and transferrin receptor 1 (TFRC), which has been shown to be cleaved by TMPRSS6^29^. Of the remaining four candidate proteins, β-Klotho (KLB) and Insulin Receptor (INSR) have both been implicated in hepatic lipid metabolism and MASLD^30–33^. Given the growing interest in the FGFR-KLB pathway as a therapeutic target for MASLD^34–36^, KLB was selected for further investigation.

**Table 1.** Proteins significantly enriched in TMPRSS6 WT compared with both Mock and TMPRSS6 S762A conditions.

| Gene | Protein | Subcellular location | Liver RNA expression (nTPM)<sup>26</sup> | Function |
| --- | --- | --- | --- | --- |
| CA12 | Carbonic anhydrase 12 | Plasma membrane | 0.7 | pH regulation |
| F11 | Coagulation factor XI | Secreted | 227.2 | Coagulation pathway |
| F11R | Junctional adhesion molecule A | Plasma membrane | 68.8 | Tight junction formation |
| INSR | Insulin receptor | Plasma membrane | 60.3 | Insulin signaling |
| ITM2C | Integral membrane protein 2C | Plasma membrane | 35.1 | APP regulation |
| KLB | Beta-klotho | Plasma membrane | 16.6 | FGF signaling |
| TFRC | Transferrin receptor protein 1 | Plasma membrane | 26.1 | Cellular iron uptake |
| TMPRSS6 | Transmembrane protease serine 6 | Plasma membrane | 129.5 | Iron and lipid metabolism |

### TMPRSS6 interacts with and cleaves KLB

KLB functions as the obligate co-receptor for FGF21 and FGF19 signaling^37–39^. Whereas FGF21 primarily activates FGFR1c and FGFR3c^40^, FGF19 preferentially signals through FGFR4^37^. In the liver, FGFR4 is the predominant FGF receptor^41^ and activation of the FGF19-FGFR4-KLB signaling axis promotes beneficial metabolic effects by reducing bile acid production^42^, gluconeogenesis^43^ and lipogenesis^44^ while increasing fatty acid oxidation^45^ (Fig. 2A). Therefore, proteolytic cleavage of the KLB ectodomain would be expected to impair FGF19 signaling, providing a potential mechanism through which TMPRSS6 may contribute to MASLD pathogenesis. Structurally, KLB is a 1044 amino acid type I transmembrane protein with a short cytoplasmic domain and a large ectodomain composed of two glycoside hydrolase-like domains (KL1 and KL2)^46,47^. Although the proteolytic processing of KLB remains poorly characterized, it has been proposed by analogy with α-klotho (KL) that KLB undergoes cleavage by a disintegrin and metalloproteinase 10 (ADAM10) and 17 (ADAM17) between the KL2 domain and transmembrane domain (α-cut) or between KL1 and KL2 domains (β-cut)^31,48,49^ (Fig. 2B).

**Figure 2.**
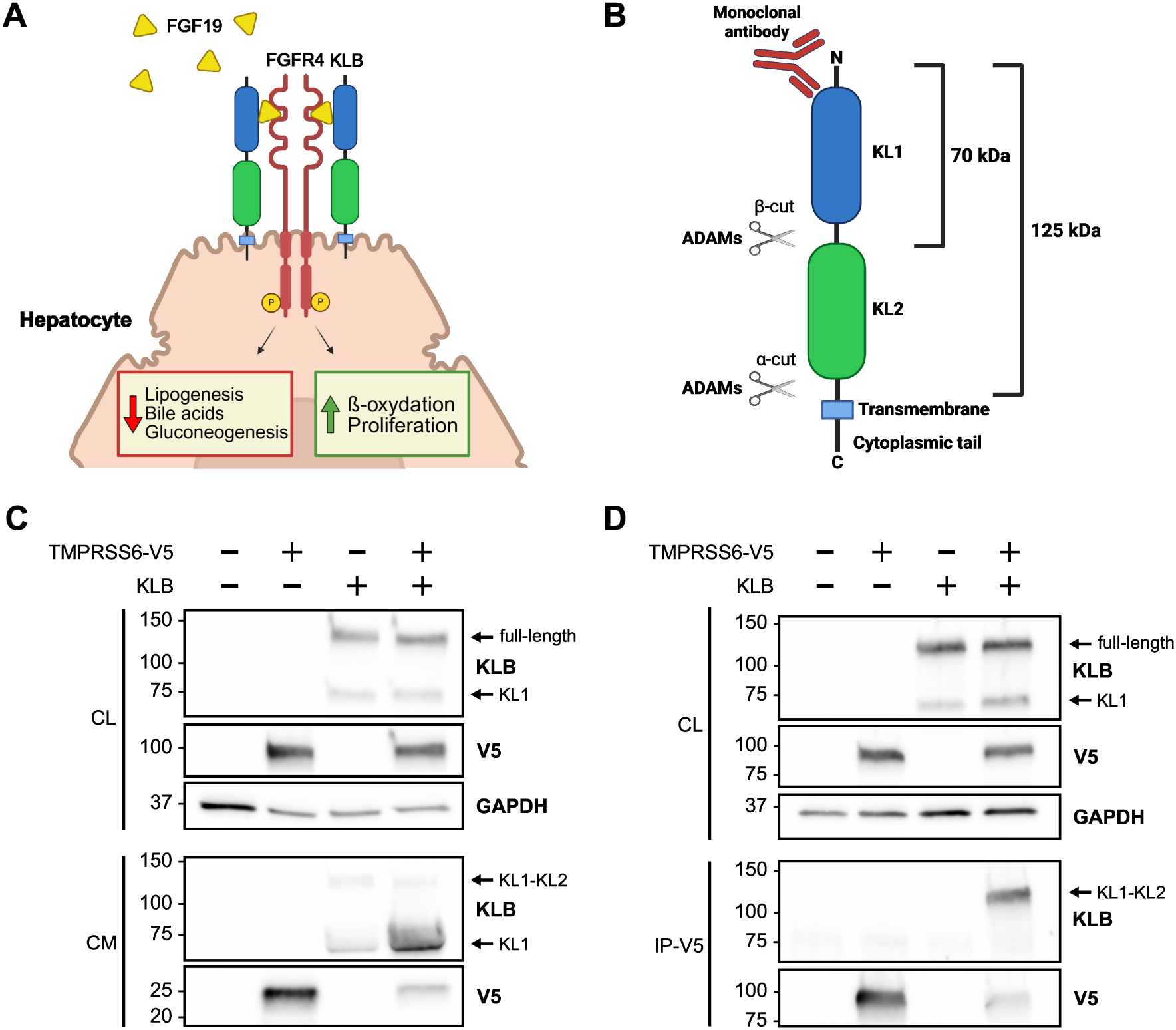
TMPRSS6 interacts with and cleaves KLB. (A) Role of KLB in the liver. KLB functions as the obligate co-receptor of FGFR4, enabling its activation by FGF19. In hepatocytes, activation of the FGF19-FGFR4-KLB signaling axis decreases lipogenesis, bile acid production and gluconeogenesis while increasing β-oxidation and cell proliferation. (B) Structure of KLB. KLB is a type I transmembrane protein composed of two inactive glycosyl hydrolase-like domains (KL1 and KL2), a single transmembrane domain and a short C-terminal cytoplasmic tail. Cleavage by ADAM10/17 generates a 125 kDa fragment (α-cut) or a 70 kDa fragment (β-cut). The monoclonal antibody used for subsequent immunoblotting recognizes the N-terminal region of the ectodomain. (C) TMPRSS6-V5 and KLB were transfected alone or together in HEK293SL cells. Cell lysates (CL) and cell media (CM) were collected and analyzed by immunoblotting using the indicated antibodies. GAPDH was used as a loading control for cell lysates. (D) TMPRSS6-V5 and KLB were transfected alone or together in HEK293SL cells. Whole-cell lysates were immunoprecipitated using anti-V5 magnetic beads. Cell lysates (CL) and immunoprecipitated proteins (IP-V5) were analyzed by immunoblotting using the indicated antibodies. GAPDH was used as a loading control for cell lysates.

To further characterize KLB as a potential substrate of TMPRSS6, we first investigated whether TMPRSS6 promotes KLB shedding. Lysates and conditioned media from HEK293SL cells overexpressing KLB, TMPRSS6 or both were analyzed by immunoblotting (Fig. 2C). In cell lysates, two forms of KLB were detected, full-length KLB and a 70 kDa fragment consistent with the released KL1-domain following β-cut cleavage. In conditioned medium, expression of KLB alone resulted in the shedding of two extracellular fragments corresponding to the full ectodomain generated by α-cut cleavage and the KL1-containing fragment generated by β-cut cleavage, indicating endogenous KLB shedding in HEK293SL cells. Importantly, co-expression of TMPRSS6 and KLB markedly increased the abundance of the 70 kDa fragment in the conditioned medium (Fig. 2C), demonstrating that TMPRSS6 promotes KLB shedding. Next, to determine if TMPRSS6 directly cleaves KLB or promotes its shedding through an indirect mechanism, we investigated whether the two proteins interact. V5-tagged TMPRSS6 and KLB were co-transfected in HEK293SL cells followed by immunoprecipitation of V5-tagged TMPRSS6. Cell lysates and immunoprecipitated proteins were then analyzed by western blot (Fig. 2D). KLB was detected in V5-immunoprecipitated proteins only when co-expressed with V5-TMPRSS6, indicating an interaction between the proteins (Fig. 2D).

### TMPRSS6-mediated KLB shedding requires catalytic activity

To assess whether KLB shedding is specifically mediated by catalytically active TMPRSS6, we compared wild-type TMPRSS6 with inactive TMPRSS6 and other members of the TTSP family (Fig. 3A). HEK293SL cells were co-transfected with KLB along with wild-type TMPRSS6, the inactive mutant TMPRSS6 S762A, the inactive isoform 4 of TMPRSS6^29^, hepsin or matriptase. Hepsin is the other major TTSP expressed in the liver^50^, whereas matriptase is a closely related TTSP having high homology with TMPRSS6 and found in most epithelial tissues^51^. Only the catalytically active TMPRSS6 WT increased the abundance of the 70 kDa fragment in the conditioned medium. Neither TMPRSS6 S762A nor TMPRSS6 isoform 4 affected KLB shedding compared with KLB expressed alone. Co-expression of hepsin or matriptase did not increase the 70 kDa KLB fragment in the conditioned media but led to a drastic reduction of KLB in cell lysates, suggesting non-specific degradation rather than specific ectodomain shedding, possibly reflecting overexpression artifacts (Fig. 3A).

We next investigated whether TMPRSS6-mediated cleavage of KLB could be modulated by endogenous or exogenous TMPRSS6 inhibitors. To this end, HEK293SL cells were co-transfected with TMPRSS6 WT, KLB and hepatocyte growth factor activator inhibitor type 2 (HAI2), a Kunitz-type serine protease that has been previously shown to inhibit TMPRSS6^52,53^. In parallel, cells co-expressing TMPRSS6 WT and KLB were treated with Compound-8 (Cpd8), a potent peptidomimetic TMPRSS6 inhibitor previously developed by our group^54,55^. Both HAI-2 and Cpd-8 completely abolished TMPRSS6-dependent KLB shedding (Fig. 3B), further supporting that the increased shedding of KLB requires TMPRSS6 proteolytic activity.

**Figure 3.**
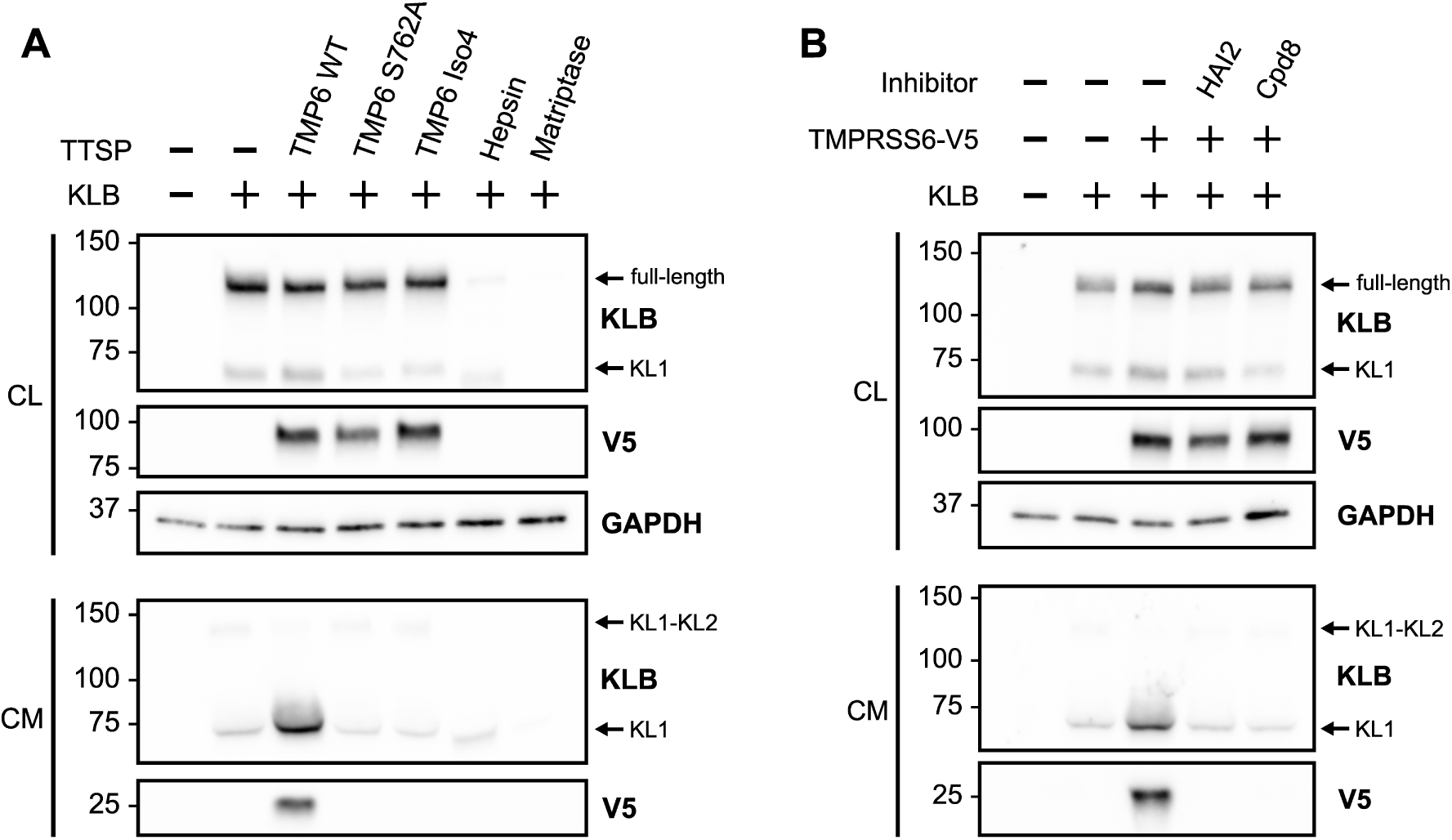
TMPRSS6-mediated KLB shedding requires catalytic activity. (A) HEK293SL cells were co-transfected with KLB together with wild-type V5-TMPRSS6 (TMP6 WT), the catalytically inactive V5-TMPRSS6-S762A mutant (TMP6 S762A), V5-TMPRSS6 isoform 4 (TMP6 iso4), hepsin, or matriptase. Cell lysates (CL) and conditioned media (CM) were collected and analyzed by immunoblotting using the indicated antibodies. GAPDH was used as a loading control for cell lysates. (B) HEK293SL cells were co-transfected with TMPRSS6-V5 and KLB with an empty vector or hepatocyte growth factor-activator activator inhibitor type 2 (HAI2). Cells were treated overnight with DMSO 0,1% or 10µM of compound 8 (Cpd8). Cell lysates (CL) and conditioned media (CM) were collected and analyzed by immunoblotting using the indicated antibodies. GAPDH was used as a loading control for cell lysates.

### TMPRSS6 mediates KLB shedding despite ADAM inhibition

Because the KLB fragment that is increasingly shed in presence of TMPRSS6 is also generated endogenously in HEK293SL cells, we investigated whether TMPRSS6 promotes KLB shedding indirectly through ADAM activation or by directly cleaving KLB. HEK293SL cells were transfected with KLB alone or co-transfected with TMPRSS6, ADAM10 or ADAM17. Cells were then treated with either Cpd8, a TMPRSS6 inhibitor, or TAPI-1, a broad-spectrum ADAM and matrix-metalloproteinase (MMP) inhibitor previously shown to block α-cut and β-cut cleavage of α-klotho^49^. Cell lysates and conditioned media were analyzed by immunoblotting.

**Figure 4.**
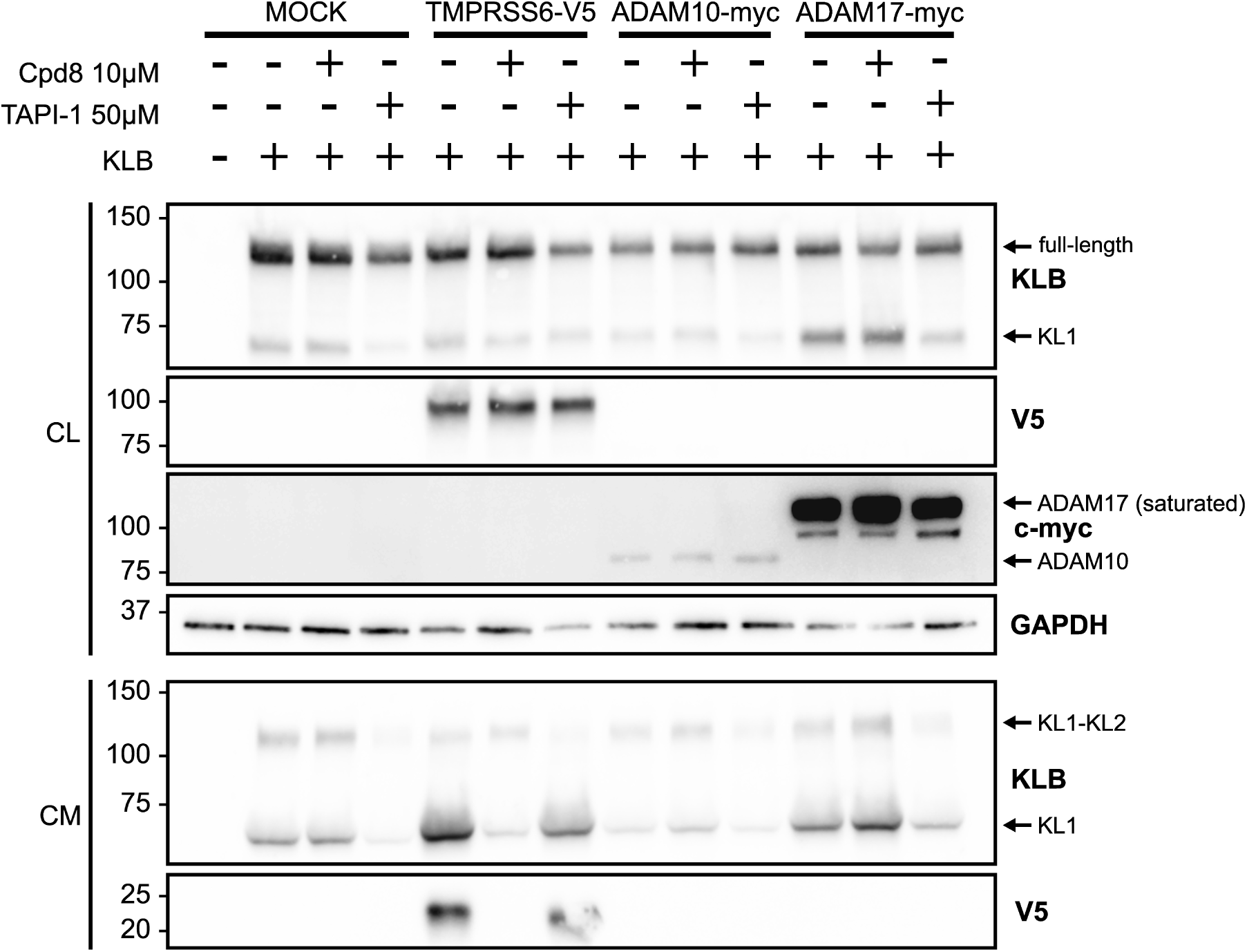
TMPRSS6 mediates KLB shedding despite ADAM inhibition. HEK293SL cells were co-transfected with KLB along with an empty vector, TMPRSS6-V5, ADAM10 or ADAM17. Cells were treated overnight with vehicle (DMSO 0,1%) or 10µM of compound 8 (Cpd8) or 20 to 50µM of TAPI-1. Cell lysate (CL) and conditioned media (CM) were collected and analyzed by immunoblotting using the indicated antibodies. V5 and c-myc were used as expression controls for TMPRSS6 and ADAM10/17, respectively. GAPDH was used as a loading control for cell lysates.

When KLB is expressed alone, Cpd8 had no effect on basal KLB shedding while TAPI-1 markedly reduces both the full shed ectodomain and KL1 fragment generated respectively by the α-cut and β-cut. Conversely, the marked increase in KL1 fragment shedding induced by TMPRSS6 was completely abolished by Cpd8 but only partially reduced by TAPI-1, indicating that TMPRSS6 can promote KLB shedding independently of ADAM activity. Co-expression of KLB with ADAM10 produced a shedding pattern comparable to that of KLB alone, suggesting that ADAM10 does not contribute to KLB processing in this model. By contrast, ADAM17 increased shedding of the KL1 fragment, although to a lesser extent than TMPRSS6. This increase was not affected by Cpd8 but was completely blocked by TAPI-1. Taken together, these findings suggest that ADAM17 but not ADAM10 contributes to endogenous KLB shedding in HEK293SL cells, whereas TMPRSS6 is able to promote KLB shedding even when ADAM activity is inhibited.

### Identification of the TMPRSS6 cleavage sites in KLB

In the previous experiments, we used a monoclonal anti-KLB antibody recognizing the N-terminal region of KLB (Fig. 2B). To facilitate the identification of TMPRSS6 cleavage sites within KLB, we next analyzed conditioned media from HEK293SL cells co-expressing KLB and TMPRSS6 using a polyclonal anti-KLB antibody, allowing detection of fragments derived from different regions of the protein. This antibody indeed detected multiple shed fragments of KLB (Fig. 5A). In addition to the previously observed 70 kDa fragment using the monoclonal antibody, another major fragment of approximately 37 kDa and a minor fragment of 55 kDa were detected (Fig. 5A). To identify the regions contained within each fragment and determine potential TMPRSS6 cleavage sites, the three KLB fragments were isolated and digested either with chymotrypsin or LysC, proteases cleaving respectively after aromatic/hydrophobic residues^56^ or after lysine residues^57^. Since TMPRSS6 is known to preferentially cleave after arginine residues^58^, this method was designed to identify peptide sequences compatible with TMPRSS6-mediated cleavage. The resulting peptides were then analyzed by mass spectrometry (schematic representation of identified fragments in Fig. 5B). Peptides ranging from residues N84 to Y551 were identified in the 70 kDa fragment (yellow/orange) indicating that it contains the KL1 domain along with the N-terminal portion of the KL2 domain. The 55 kDa fragment (red/orange), which was less abundant, contained peptides extending from N84 to F455 suggesting the presence of a less prominent cleavage site located downstream of F455. Finally, peptides identified in the 37 kDa fragment (green) comprised residues from T571 to R855, which correspond to most of the KL2 domain. Together, these data indicate that the cleavage site generating the 70 kDa fragment is located between amino acids Y551 and T571 in the KL2 domain. For the 37 kDa fragment, R855 emerged as a particularly compelling candidate cleavage site because the C-terminal arginine residue R855 cannot be generated by either chymotrypsin or LysC digestion during sample preparation and it corresponds to the preferred cleavage residue of TMPRSS6 (Fig. 5B).

**Figure 5.**
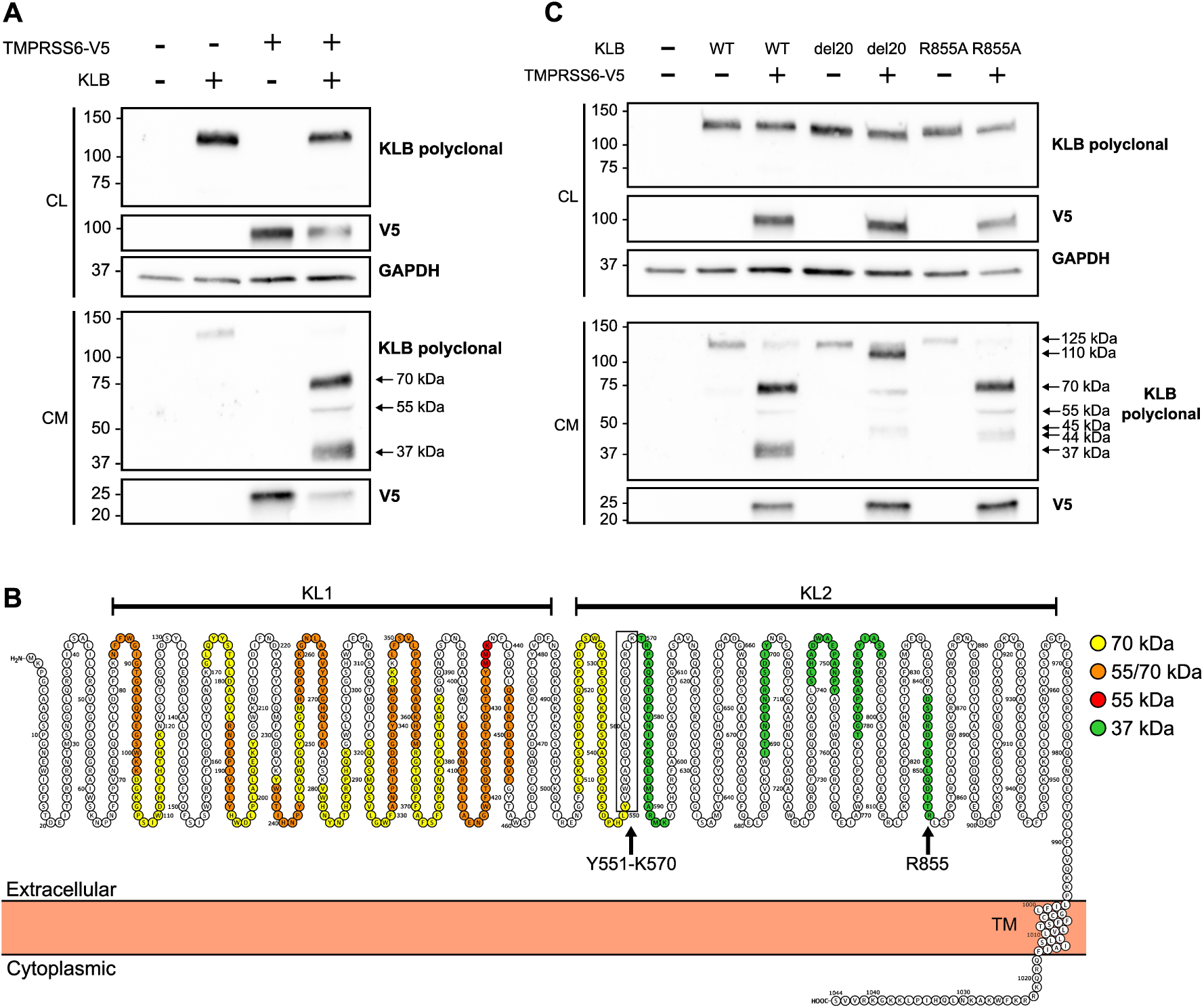
Identification of the TMPRSS6 cleavage sites in KLB. (A) TMPRSS6-V5 and KLB were transfected alone or together in HEK293SL cells. Cell lysate (CL) and cell media (CM) were collected and analyzed by immunoblotting using the indicated antibodies. GAPDH was used as a loading control for cell lysates. (B) Snake plot of KLB showing the peptides identified by mass spectrometry in the shed KLB fragments detected in (A). Excised protein bands were digested with either chymotrypsin or LysC prior to mass spectrometry analysis. Amino acids identified only in the 70 kDa fragment are shown in yellow, those identified in both the 70 and 55 kDa fragments in orange, those identified exclusively in the 55 kDa fragment in red and those identified in the 37 kDa fragment in green. Based on these peptides distribution, two potential cleavage regions were identified, the region between Tyr551 and Lys570 (Y551-K570) and Arg855 (R855). (C) Cleavage of KLB mutants. HEK293SL cells were co-transfected with TMPRSS6-V5 and either wild-type KLB (WT), a KLB mutant with a 20 amino acids deletion from Y551 to K570 (del20), or a KLB mutant containing R855A substitution. Cell lysate (CL) and cell media (CM) were collected and analyzed by immunoblotting using the indicated antibodies. GAPDH was used as a loading control for cell lysates.

Given the low abundance of the 55 kDa fragment and the inability to localize its cleavage to a narrow region, we focused subsequent analyses on the major 70 and 37 kDa cleavage products. To validate the candidate cleavage regions, two KLB mutants were generated: one carrying a deletion of 20 amino acids spanning Y551 to K570 (del20) and another harboring an arginine to alanine substitution at position 855 (R855A). These mutants were then expressed in HEK293SL cells with or without TMPRSS6 and conditioned media were analyzed by immunoblotting using the polyclonal anti-KLB antibody (Fig. 5C, Fig. S2 for overexposed membrane). As anticipated, co-expression of TMPRSS6 with the KLB del20 mutant markedly reduced the abundance of the 70 and 37 kDa fragments. Instead, two new fragments of approximately 110 and 45 kDa appeared, consistent with loss of cleavage in the Y551-K570 region. The 110 kDa fragment is consistent with the ectodomain extending from the N-terminal to R855, and the 45 kDa fragment likely represents the C-terminal portion of this last fragment after the alternative cleavage downstream of F455. Interestingly, in absence of TMPRSS6, the KLB del20 mutant also blocked the endogenous shedding of the 70 kDa fragment (Fig. S2), indicating that deletion of Y551-K570 interferes with β-cut. In contrast, co-expression of TMPRSS6 with the KLB R855A mutant resulted in unchanged shedding of the 70 kDa fragment and the complete loss of the 37 kDa band, supporting R855 as a TMPRSS6 cleavage site. As expected, a new fragment of approximately 44 kDa was detected, consistent with the region extending from the α-cut site to the cleavage site within the Y551-K570 region (Fig. 5C). Altogether, these results indicate that TMPRSS6 facilitates cleavage in the Y551-K570 region and specifically cleaves at the R855 residue in the KL2 domain.

### TMPRSS6 reduces full-length KLB at the cell surface and impairs FGF19-mediated FGFR4 signaling

Because KLB functions as a cell-surface co-receptor, its expression at the plasma membrane is required to support cellular signaling. Although TMPRSS6 increased KLB shedding into the conditioned medium, we did not observe a corresponding decrease in full-length KLB in cell lysates (Fig. 1C). We therefore investigated whether TMPRSS6-mediated cleavage was sufficient to reduce KLB cell-surface expression. HEK293SL cells transfected with KLB, TMPRSS6, or both were subjected to cell-surface biotinylation, and biotinylated proteins were analyzed by immunoblotting (Fig. 6). When KLB was expressed alone, both full-length KLB (125 kDa) and the 70 kDa fragment were detected in whole cell lysates and in the biotinylated cell-surface proteins. In contrast, co-expression of TMPRSS6 markedly reduced the abundance of cell-surface full-length KLB (125 kDa) fraction to near-undetectable levels, whereas the reduction of the 70 kDa fragment was considerably less pronounced. Expression of full-length KLB in the cell lysate was not affected. These results indicate that TMPRSS6-dependent cleavage reduces the amount of full-length KLB present at the cell surface, potentially impairing its function as a co-receptor.

**Figure 6.**
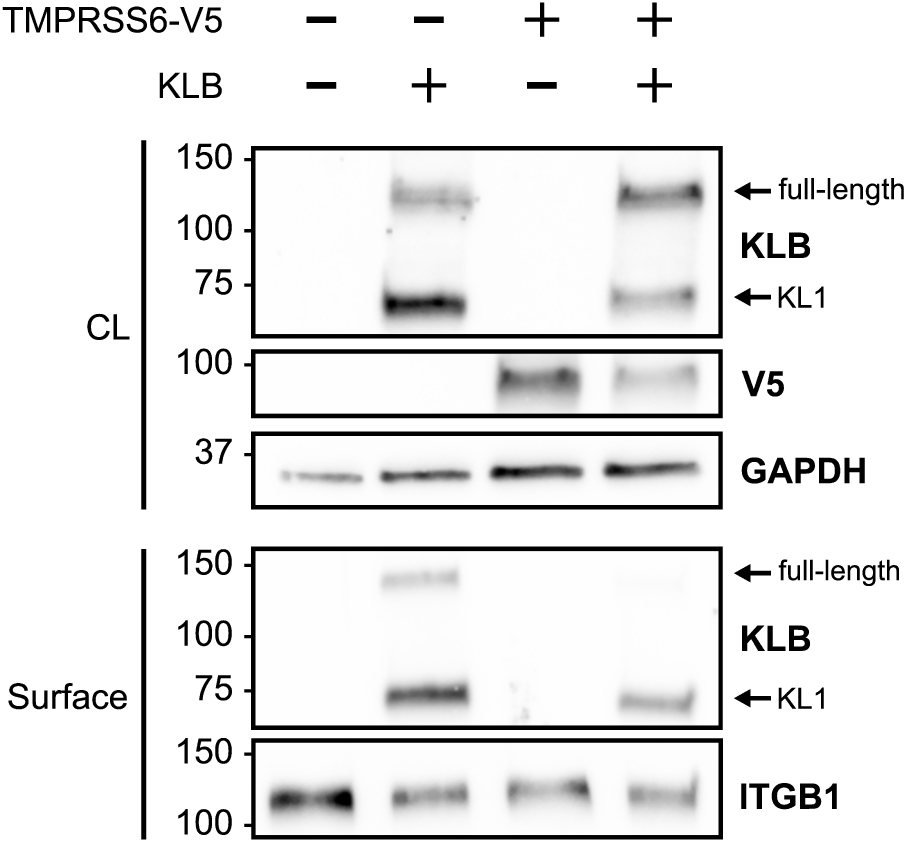
TMPRSS6 reduces full-length KLB at the cell surface. TMPRSS6-V5 and KLB were transfected alone or together in HEK293SL cells. After a 24-h transfection, cell surface proteins were biotinylated and isolated by avidin precipitation. Cell lysates (CL) and biotinylated cell surface proteins (Surface) were analyzed by immunoblotting using the indicated antibodies. GAPDH was used as a loading control for cell lysates and ITGB1 as a loading control for cell surface proteins.

Because KLB functions as the obligate co-receptor for FGF19 and FGF21^37–39^, we investigated whether TMPRSS6-mediated KLB cleavage affects FGF19-dependent activation of FGFR4, the main FGF receptor in hepatocytes^41^. HEK293SL cells were transfected with KLB and FGFR4, either alone or with TMPRSS6 WT or the catalytically inactive TMPRSS6 S762A mutant. Following FGF19 stimulation, activation of the mitogen-activated protein kinase (MAPK) pathway, a major downstream effector of FGFR4 signaling^59,60^, was assessed using dual-luciferase reporter assay. Firefly luciferase with the serum response element (SRE) promoter was used to measure MAPK activity, and Renilla Luciferase driven by the simian virus 40 (SV40) promoter served as a transfection efficacy control (Fig. 7). As expected FGF19 failed to activate MAPK signaling in the absence of KLB and FGFR4. In contrast, co-expression of KLB and FGFR4, resulted in robust FGF19-induced MAPK activation. Interestingly, expression of wild-type TMPRSS6 modestly but significantly reduced FGF19-dependent FGFR4 signaling by approximately 20%, whereas the catalytically inactive mutant TMPRSS6 S762A showed no effect (Fig. 7).

**Figure 7.**
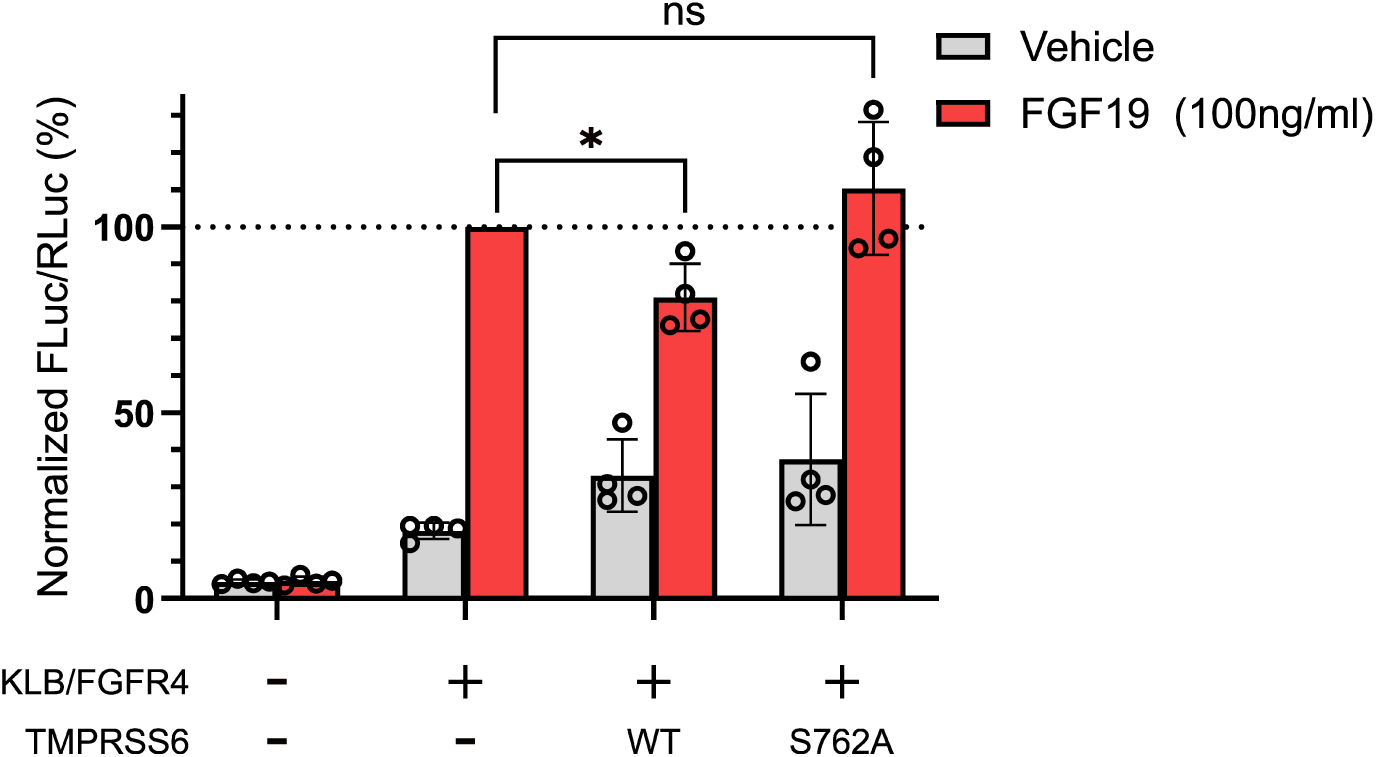
TMPRSS6 impairs FGF19-mediated FGFR4 signaling. HEK293SL cells were co-transfected with KLB and FGFR4 either alone or together with wild-type TMPRSS6 (WT) or the inactive TMPRSS6 S762A mutant (S762A). All conditions were also co-transfected with the SRE firefly luciferase reporter and the SV40 Renilla luciferase normalization plasmid. Twenty-four hours after transfection, cells were serum-starved overnight and stimulated with FGF19 or vehicle for 6 hours. Firefly and Renilla luciferase activities were measured sequentially using the Dual-Glo Luciferase Reporter Assay System. Relative MAPK pathway activation was determined by normalizing firefly luciferase activity to the corresponding *Renilla* luciferase activity. Values were then normalized to the KLB/FGFR4 without TMPRSS6 condition. Statistical significance was assessed using a one-sample t-test comparing TMPRSS6 WT and TMPRSS6 S762A to the control value of 100%.

These findings indicate that TMPRSS6 proteolytic activity attenuates FGF19 signaling in this model, consistent with its ability to reduce cell-surface KLB expression. Taken together, our data identify KLB as a functional substrate of TMPRSS6 and demonstrate that its cleavage attenuates FGF19/FGFR4 signaling in an overexpression system.

## Discussion

Because of its key role in iron homeostasis, TMPRSS6 has emerged as an attractive therapeutic target for several iron overload disorders such as hemochromatosis, β-thalassemia, and polycythemia vera^61–63^. Recently, growing evidence has also implicated TMPRSS6 in hepatic lipid metabolism and MASLD^20–22^. These findings raise the possibility that TMPRSS6 could represent a therapeutic target beyond iron disorders and highlight the importance of better understanding the molecular mechanisms underlying its role in metabolic liver disease.

In the present study, we used an extracellular proteomic approach to identify novel potential substrates of TMPRSS6. Importantly, TFRC, which has already been shown to be cleaved by TMPRSS6^29^, was identified, supporting the validity of the experimental approach. Among the newly identified candidates, insulin receptor (INSR) also emerged as a particularly interesting hit. Indeed, previous studies have shown that the insulin receptor can be proteolytically cleaved in the liver by calpain-2 or BACE1, resulting in impaired insulin signaling^64,65^. Given the central role of insulin resistance in hepatic lipid metabolism and MASLD,^33,66,67^ it will be important to determine whether TMPRSS6 similarly processes INSR. Nonetheless, KLB was selected for further investigation because of its well-established role in MASLD^30,31,68^. KLB functions as the obligate co-receptor for FGF19 and FGF21 signaling^37–39^. Agonists for both FGF19 and FGF21 are currently being developed as therapies for MASH and have demonstrated promising effect in clinical studies by increasing fatty acid oxidation, reducing fibrosis and improving resolution of MASH in clinical studies^34–36^. Therefore, proteolytic cleavage of KLB, which would be expected to impair FGF19 and FGF21 signaling in the liver, represents a plausible mechanism by which TMPRSS6 could contribute to MASLD.

Proteolytic shedding of KLB has previously been hypothesized to be mediated by ADAM10 and ADAM17, largely by analogy with α-klotho (KL)^31,48,49^. Our findings identify TMPRSS6 as an additional protease capable of processing KLB. Importantly, the increased KLB shedding induced by TMPRSS6 was abolished by the endogenous inhibitor HAI2^52^ and a selective peptidomimetic TMPRSS6 inhibitor^54,55^, demonstrating that this cleavage depends on TMPRSS6 catalytic activity. Moreover, neither the closely related matriptase nor the liver TTSP hepsin, reproduced the characteristic cleavage pattern observed with TMPRSS6. In addition, TMPRSS6 retained the ability to promote KLB shedding in the presence of the broad-spectrum ADAM inhibitor TAPI-1, indicating that its effect cannot be explained solely by activation of endogenous ADAM proteases. Instead, these findings support a direct proteolytic role for TMPRSS6 in KLB processing.

Mass spectrometry analysis of shed KLB fragments identified two potential TMPRSS6 cleavage regions that were subsequently analyzed using KLB mutants: the region from Tyr551 to Lys570 and Arg855. While deletion of residues Y551-K570 markedly reduced generation of the TMPRSS6-dependent fragments, it also impaired endogenous KLB shedding, indicating that this region likely contains the β-cut cleavage site of the ADAM sheddases. This observation suggests that, unlike KL^69^, the β-cut cleavage site of KLB is not located in the unstructured linker between the KL1 and KL2 domains but rather within the N-terminal portion of KL2, in a loop known in KL as the receptor binding arm (RBA)^70^. In contrast, substitution of Arg855 by alanine abolished generation of the 37 kDa fragment without affecting basal shedding. Together with the known preference of TMPRSS6 for cleavage after arginine residues, these findings strongly support Arg855 as a direct TMPRSS6 cleavage site.

Although TMPRSS6 clearly promoted KLB shedding, total KLB abundance in whole-cell lysates remained unchanged. This prompted us to investigate whether TMPRSS6-mediated processing affects the amount of functional KLB at the plasma membrane. Cell-surface biotinylation showed that TMPRSS6 reduces the full-length form of KLB at the cell surface. Consistent with this observation, TMPRSS6 attenuated FGF19-dependent activation of FGFR4 in an overexpression system. Given that FGFR4 is the main FGF receptor in hepatocytes, these findings uncover a potential mechanism by which TMPRSS6 may contribute to hepatic steatosis, although validation in hepatocytes and *in vivo* models will be required to establish physiological relevance.

By cleaving KLB, TMPRSS6 would be expected to impair endocrine FGF signaling, thereby limiting the metabolic benefits of both FGF19 and FGF21. FGF19 primarily acts on the liver to reduce bile acid production^42^, gluconeogenesis^43^ and lipogenesis^44^ and stimulate β-oxidation^45^ and glycogen synthesis^71^ but is also has been shown to increase production of FGF21 by the liver^72^. FGF21 acts on adipose tissue to stimulate lipolysis and improve insulin sensitivity via adiponectin production^73^, on the liver to reduce hepatic cholesterol^74^ and on the brain by promoting a preference for protein intake over carbohydrate and fat consumption^75^. Interestingly, FGF21 expression is also regulated by PPARα^76^, raising the possibility of an intersection between FGF signaling and the previously proposed BMP-SMAD-PPARα mechanism of TMPRSS6 action^21^. In addition, it will be important to determine the impact of TMPRSS6-mediated KLB cleavage on cell proliferation and tumorigenesis, as FGF19-FGFR4 signaling has been implicated in the development of hepatocellular carcinoma^77–80^. It would also be interesting to investigate the potential biological role of the shed 70 kDa KLB fragment, as the circulating soluble ectodomain of the related protein KL has been shown to exert endocrine effects^70,81–83^.

From a therapeutic perspective, our findings further support the rationale for targeting TMPRSS6 in MASLD. TMPRSS6 has already emerged as a promising therapeutic target for iron-overload disease as several molecular tools are in development. Multiple therapeutic strategies including antisense oligonucleotides (ASO), small interfering RNA (siRNA) and antibody targeting TMPRSS6 have shown efficacy in preclinical models and early-phase clinical trials by increasing hepcidin, reducing iron tissue overload, and improving hematological parameters^84–90^. Our study suggests that, beyond its established role in iron homeostasis, inhibition of TMPRSS6 may also preserve KLB-dependent FGF signaling, providing additional justification for investigating TMPRSS6-targeted therapies in MASLD.

Collectively, our findings identify KLB as a novel functional substrate of TMPRSS6 and show that its proteolytic processing reduces cell-surface KLB abundance and attenuates FGF19/FGFR4 signaling in a heterologous expression system. By linking TMPRSS6 activity to a key metabolic signaling pathway, our findings provide a mechanistic framework through which TMPRSS6 may contribute to MASLD and support continued investigation of TMPRSS6 as a therapeutic target in metabolic liver disease. Future studies in physiological models will be required to establish the *in vivo* relevance of TMPRSS6-mediated KLB cleavage and to further define the contribution of TMPRSS6 catalytic activity to its diverse biological functions.

## Material and Methods

### Cell Culture, Reagents, and Antibodies

Human HEK293SL (from Stephane Laporte laboratory, McGill, Canada), a 293-cell line subclone selected for enhanced adherence^91^, were maintained in Dulbecco’s Modified Eagle’s Medium High Glucose (DMEM) (Wisent, 319-005-CL). Hep3B cells were purchased from American Type Culture Collection (ATCC) and cultured in Eagle’s Minimum Essential Medium (EMEM) (Wisent, 320-005-CL). Both media were supplemented with 10% Fetal Bovine Serum (FBS; Wisent, 080-150) and 1% antibiotic mixture (penicillin-streptomycin-L-glutamine; Wisent, 450-202-EL). HCELL-100 (001-035-CL) and D-PBS (311-425-CL) were acquired from Wisent. Opti-MEM (31985062) was purchased from Gibco and lipofectamine 3000 (L3000015) from Invitrogen. 10 kDa Amicon centrifugal filters (UFC501096) were purchased from Merck Millipore. Anti-V5 magnetic beads (M167-11) were acquired from MBL Life science. Pierce Cell Surface Biotinylation and Isolation Kit (A44390) was obtained from Thermo Fisher Scientific. Dual-Glo Luciferase Assay System (E2920) was purchased from Promega.

Mouse monoclonal anti-V5 antibody (R96025) was purchased from Invitrogen. Monoclonal mouse anti-KLB (MAB5889) and polyclonal goat anti-KLB (AF5889) were obtained from R&D systems. HRP-linked rabbit anti-GAPDH (8884S), HRP-linked mouse anti-Myc (2040S) and rabbit anti-ITGB1 (4706S) were purchased from Cell Signaling Technology. Rabbit anti-goat IgG secondary antibody (A16136) was purchased from Invitrogen. Horse anti-mouse IgG secondary antibody (7076S) and goat anti-rabbit IgG secondary antibody (7074S) was obtained from Cell Signaling Technology.

### Plasmid construction

Construct encoding TMPRSS6-V5 WT (isoform 2), TMPRSS6-V5 S762A (isoform 2) and TMPRSS6 isoform 4 were obtained as previously described^92,93^. Matriptase construct was obtained as previously described^94^. KLB WT coding sequence was synthesized and subcloned in pcDNA 3.1 (+) vector (Bio Basic). KLB mutant constructs were obtained using the QuikChange site-directed mutagenesis kit (Agilent Technologies). C-terminal HA-tagged Hepsin construct was subcloned in pcDNA 3.1(+) (GenScript). HAI-2 construct was a gift from Karin List (Wayne State University, USA). ADAM10-myc and ADAM17-myc were a gift from Renata Mężyk-Kopeć (Uniwersytet Jagielloński, Poland). pcDNA3.1-FGFR4-V5/HIS (PMID28199182) was a gift from Pavel Krejčí (Addgene plasmid # 201109; http://n2t.net/addgene:201109 ; RRID:Addgene_201109). Plasmids used in luciferase assays (pGL4.33 (luc2P/SRE/Hygro) and pGL4-73 (hRluc/SV40)) are from Promega.

### Transfection of TMPRSS6 in Hep3B cells

Hep3B cells (5 x 10^5^) were seeded in 60 mm dishes (Corning) and allowed to grow overnight. The next day, cells were transfected with 1.875 µg of a plasmid encoding TMPRSS6-V5, TMPRSS6-V5 S762A or pcDNA3.1 using Lipofectamine 3000 according to the manufacturer’s instructions. 24 hours post-transfection, cells were washed with D-PBS and cell media was changed for serum-free EMEM. Cells were incubated for 48 hours after which conditioned media were collected for downstream protein analysis.

### Precipitation and digestion of extracellular proteins

Proteins present in conditioned media were concentrated by TCA/acetone precipitation. Briefly, one volume of 20% TCA/acetone solution was added to one volume of conditioned media. Samples were mixed and incubated on ice for 5 minutes. Samples were then centrifuged at 15,000g for 3 minutes. Protein pellets were washed with ice-cold acetone, followed by centrifugation at 15 000g for 2 minutes. This washing step was repeated twice using 80% ice-cold acetone. Pellets were air-dried and resuspended in lysis buffer (5% SDS, 50mM TEAB, pH 8.5).

Equal amounts of proteins were processed using S-Trap micro spin columns (Protifi) according to the manufacturer’s instructions. Proteins were reduced with 5 mM of tris(2-carboxyethyl)phosphine (TCEP) for 15 min, alkylated with 20 mM of chloroacetamide (ClAA) for 10 min, and acidified with 2.5% phosphoric acid. Proteins were trapped on the S-Trap matrix, digested with trypsin overnight, and peptides were eluted sequentially with 50 mM triethylammonium bicarbonate (TEAB), 0.2% formic acid and 50% acetonitrile. Pooled eluates were dried and resuspended in 1% formic acid.

### Mass spectrometry analysis in diaPASEF mode

Mass spectrometry analysis of peptides was carried out by the Université de Sherbrooke Proteomic Platform. Digested peptides (375 ng) were injected into an HPLC (nanoElute, Bruker Daltonics) equipped with a trap column (Acclaim PepMap100 C18, 0.3 mm id × 5 mm, Dionex Corporation) and analytical C18 column (1.9 μm beads, 75 μm × 25 cm, PepSep, Bruker Daltonics). Peptides were separated over 2 h using a linear gradient of 5–37% CH3CN in 0.1% FA at a flow rate of 400 nL·min−1 while being injected into a TimsTOF Pro ion mobility mass spectrometer equipped with a Captive Spray nano electrospray source (Bruker Daltonics). The target intensity was set to 20 000, with an intensity threshold of 2500. Samples processing was acquired using diaPASEF mode. Briefly, for each single TIMS (100 ms) in diaPASEF mode, we used 1 mobility window consisting of 27 mass steps (m/z between 114 to 1414 with a mass width of 50 Da) per cycle (1.27 s duty cycle) and collision energy of 42.0 eV. These steps cover the diagonal scan line for +2 and +3 charged peptides in the m/z-ion mobility plane.

### Protein identification with DIA-NN

The DIA raw files were analyzed using DIA-NN software (version 1.9.2) and the Uniprot human proteome database (10/03/2024, 54,825 entries), as described previously^95^. Briefly, FASTA digest for library-free search/library generation was enabled, as well as Deep learning-based spectra, retention time and ion mobility prediction. The settings used for the analysis were: 2 miscleavages were allowed, as well as 2 variable modifications; fixed modifications were carbamidomethylation on cysteine and N-terminal methionine excision; variable modifications were methionine oxidation and N-terminal acetylation; enzyme was trypsin; peptide length between 7 and 30 amino acids; precursor charge range between 2 and 4; precursor m/z range between 100 and 1700; fragment ion m/z range between 110 to 1500; a mass tolerance of 20 ppm was fixed for both precursor ions and fragment ions, as well as a scan window of 10; precursor FDR was set to 5%; match between runs, protein inference, cross-run normalisation and peptidoforms scoring options were also enabled.

### Fold change enrichment analysis

The protein groups matrix output from DIA-NN was used for downstream analysis and keratins (KRT), cRAP proteins and protein fragments were excluded. To remove intracellular contaminant, proteins were annotated using the Gene Ontology (GO) cellular component database^96,97^ through PANTHERdb^98^. Proteins featuring the “Extracellular space”, “extracellular region”, or “External side of plasma membrane” annotations were automatically retained. Proteins annotated as “plasma membrane” were retained if Uniprot annotated them as containing a transmembrane domain. The resulting proteins dataset was analyzed with the DIA-Analyst suite from Monash University. Missing values were imputed using the Perseus-Type method, which uses random numbers drawn from a normal distribution of 1.8 standard deviation down shift and with a width of 0.3 of each sample. Fold changes and adjusted p-values (Benjamini-Hochberg method) were calculated. Proteins with |log2 fold change| ≥ 1 and adjusted p-value < 0.1 were considered significantly modulated. Results were exported and GraphPad Prism 11.0.2 was used to create volcano plots.

### Transfection in HEK293SL cells

HEK293SL cells (3 x 10^5^) were seeded on 6-well plates (Corning) and allowed to grow overnight. The next day, cells were co-transfected with 1.25 μg of each plasmid construct, or 833 ng of each plasmid when three plasmids were co-transfected. Total amount of plasmid DNA was adjusted to 2.5 µg per well with empty vector pcDNA3.1 when necessary. Transfections were performed using Lipofectamine 3000 according to the manufacturer’s instruction.

### Analysis of the conditioned media

24 hours post-transfection, cells were washed with D-PBS and cell media was changed for HCELL-100 or serum-free DMEM. The next day, 1.5 mL of cell media was collected and concentrated using 10 kDa centrifugal filter. Cell lysis was performed and equal amounts of cell lysate and volume of concentrated media were loaded on SDS-polyacrylamide gels and analyzed with immunoblotting.

### Co-immunoprecipitation of TMPRSS6 and KLB

24 hours post-transfection, cell media was refreshed. The next day, cells were washed twice with ice-cold PBS before cell lysis was performed. Equal amounts of protein were incubated overnight at 4°C with anti-V5 magnetic beads under constant rotation. Beads were subsequently washed three times with lysis buffer and bound proteins were eluted by incubation in 1x Laemmli sample buffer for 5 min at 95°C. Immunoprecipitated proteins and equal amounts of input lysates were loaded on SDS-polyacrylamide gels and analyzed with immunoblotting.

### Preparation and digestion of the gel bands containing shed KLB fragments

Bands containing the shed KLB fragments were excised from the gel, minced, washed successively with water, 50% acetonitrile, 20 mM ammonium bicarbonate, 20 mM ammonium bicarbonate/50% acetonitrile, dehydrated with 100% acetonitrile and dried. Gel pieces were reduced with 10 mM dithiothreitol (DTT) for 1h, alkylated with 50 mM ClAA for 30 min, washed with 20 mM ammonium bicarbonate and 20mM ammonium bicarbonate/50% acetonitrile, dehydrated with 100% acetonitrile and dried. In-gel digestion was done overnight using either LysC or chymotrypsin and peptides were eluted with acetonitrile and 1% formic acid. Pooled eluates were dried, resuspended in 0.1% trifluoroacetic acid and peptides were desalted using ZipTip pipet tips containing a C18 resin (Thermo Fisher Scientific). Peptides were dried and resuspended in 1% formic acid.

### Mass spectrometry analysis in PASEF mode

Mass spectrometry analysis of peptides was carried by the Université de Sherbrooke Proteomic Platform. Digested peptides (375 ng) were injected into an HPLC (nanoElute, Bruker Daltonics) equipped with a trap column (Acclaim PepMap100 C18, 0.3 mm id × 5 mm, Dionex Corporation) and analytical C18 column (1.9 μm beads, 75 μm × 25 cm, PepSep, Bruker Daltonics). Peptides were separated over 2 h using a linear gradient of 5–37% CH3CN in 0.1% FA at a flow rate of 400 nL·min−1 while being injected into a TimsTOF Pro ion mobility mass spectrometer equipped with a Captive Spray nano electrospray source (Bruker Daltonics). The target intensity was set to 20 000, with an intensity threshold of 2500. Data was acquired using data-dependent auto-MS/MS with a 100-1700 m/z mass range, with PASEF enabled with a number of PASEF scans set at 10 (1.17 s duty cycle) and a dynamic exclusion of 0.4 min, m/z dependent isolation window and collision energy of 42.0 eV. These steps cover the diagonal scan line for +2 and +3 charged peptides in the m/z-ion mobility plane.

### Peptides identification with Maxquant

The DDA raw files were analyzed using Maxquant software (version 2.6.7.0) and the Uniprot human proteome database (10/03/2024, 54,825 entries) as described previously^99^. Briefly, the settings used for the Maxquant analysis (with TIMS-DDA type in group-specific parameters) were: 4 miscleavages were allowed; fixed modifications were carbamidomethylation on cysteine and N-terminal methionine excision; variable modifications were methionine oxidation and N-terminal acetylation; enzyme was either LysC or chymotrypsin; the minimum peptide length was set to 5 amino acids; variable modifications included in the analysis were methionine oxidation and protein N-terminal. A mass tolerance of 20 ppm was used for both precursor and fragment ions. Identification values “PSM FDR”, “Protein FDR” and “Site decoy fraction” were set to 0.05. Minimum peptide count was set to 1. “Second peptides” option was also allowed. MaxQuant was run with a transfer q value of 0.3. The identified peptides were visualized as a snake plot using Protter^100^.

### Cell surface biotinylation assay

HEK293SL cells (2.5 x 10^6^) were seeded in polylysine-coated T75 flask (Corning) and allowed to grow overnight. The next day, cells were co-transfected with 10 μg of each plasmid construct and total amount of plasmid DNA was adjusted to 20 µg per flask with empty vector pcDNA3.1 when necessary. Transfections were performed using Lipofectamine 3000. After a 24-h transfection, biotinylation of HEK293SL surface proteins was performed with Pierce cell-surface protein isolation kit according to the manufacturer’s instruction. Cell lysates were precipitated (biotinylated cell-surface proteins) or not (total proteins) with avidin. The resulting samples were loaded on SDS-polyacrylamide gels and analyzed by immunoblotting.

### Luciferase reporter assays

HEK293SL cells (1.5 x 10^4^) were seeded on white 96-well plates (Corning) and allowed to grow overnight. The next day, cells were transfected with the indicated plasmids using Lipofectamine 3000. Transfection was done with 37 ng of total plasmid (15 ng of SRE-RLuc construct, 2 ng of SV40-FLuc construct, 10 ng of TMPRSS6-V5 WT or S762A, 5 ng of FGFR4-V5, 5 ng of KLB WT and pcDNA3.1 to 37ng). 24 hours post-transfection, cells were washed with D-PBS and cell media was changed for serum-free, phenol red-free DMEM. The next day, cells were treated with FGF19 100 ng/ml or vehicle for 6 hours. Luminescence of *Renilla* and firefly luciferase were sequentially measured with a Dual-Glo Luciferase Assay System following the manufacturer’s instruction. For each condition, *Renilla* luciferase luminescence reads were divided by the corresponding firefly luminescence reads. Values were then normalized to the KLB/FGFR4 without TMPRSS6 condition. GraphPad Prism 11.0.2 was used to create graphics. Statistical significance was assessed using a one-sample t-test comparing TMPRSS6 WT and TMPRSS6 S762A to the control value of 100%.

## Author contribution

Conceptualization: M.L. A.D. R.L. Analysis and interpretation of the data: M.L. A.D. G.L. Chemical synthesis M.D. Funding acquisition: P-L.B. R.L. Writing–original draft: M.L. A.D. Correction and edition of manuscript: M.L. A.D. R.L.

## Acknowledgements

We thank Renata Mężyk-Kopeć (Uniwersytet Jagielloński, Poland) for ADAM10 and ADAM17 expression vectors, Karin List (Wayne State University, USA) for HAI2 expression vector and Pavel Krejčí (Masaryk University, Czechia) for the pcDNA3.1-FGFR4-V5/HIS construct. HEK293SL cells were a kindly gift of Stéphane Laporte (McGill University, Canada). We thank Dominique Lévesque (Université de Sherbrooke Proteomic Platform) for his valuable assistance in mass spectrometry analysis. This work was supported by a grant from the Canadian Institutes of Health Research (CIHR) to RL (#PJT-175271). M.L. received research scholarships from the Fonds de Recherche du Québec-Nature et Technologie (FRQNT), the Canadian Institutes of Health Research (CIHR) and the Fonds de Recherche du Québec-Santé (FRQS). M.D. received a FRQS scholarship for this work (BF2-299838).

