## Supplementary material for "TMPRSS6 Cleavage of β-Klotho Modulates FGF19 Signaling": Suplemental files

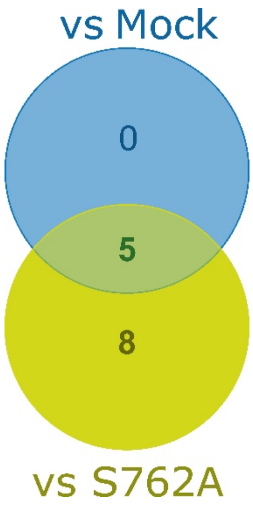


**Figure S1. Number of significantly reduced protein in TMPRSS6 WT overexpressing cells.** Venn diagram of proteins significantly reduced in the condition overexpressing TMPRSS6 WT compared to both mock and TMPRSS6 S762A.

**Table S1. Proteins significantly reduced in TMPRSS6 WT compared with both Mock and TMPRSS6 S762A conditions.**

| Gene | Protein | Subcellular location | Liver RNA expression (nTPM) | Function |
|---|---|---|---|---|
| COL6A1 | Collagen alpha-1(VI) chain | Extracellular matrix | 46.6 | Extracellular matrix scaffold |
| COMP | Cartilage oligomeric matrix protein | Extracellular matrix | 0.3 | Collagen assembly |
| HGFAC | Hepatocyte growth factor activator serine protease | Secreted | 223.6 | HGF signaling |
| ITIH3 | Inter-alpha-trypsin inhibitor heavy chain H3 | Secreted | 798.4 | Matrix stabilization |
| ITIH4 | Inter-alpha-trypsin inhibitor heavy chain H4 | Secreted | 1743.0 | Matrix stabilization |

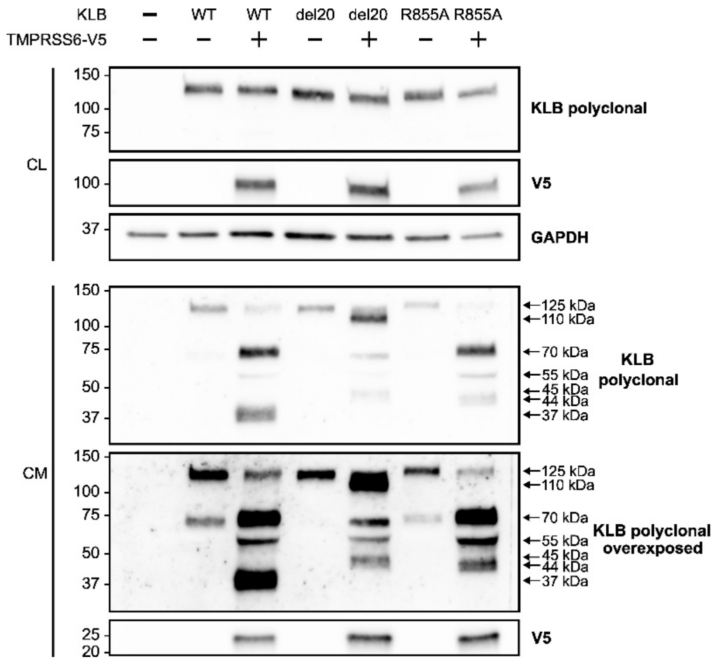


**Figure S2. Cleavage of KLB mutants.** HEK293SL cells were co-transfected with TMPRSS6-V5 and either wild-type KLB (WT), a KLB mutant with a 20 amino acids deletion from Y551 to K570 (del20), or a KLB mutant containing R855A substitution. Cell lysate (CL) and cell media (CM) were collected analyzed by immunoblotting using the indicated antibodies. An overexposed blot probed with the polyclonal KLB antibody is shown to visualize low-abundance KLB fragments. GAPDH was used as a loading control for cell lysates.
